# The Orai3 channel and AHNAK2 scaffold protein promote the activation of primary human muscle stem cells *in vitro*

**DOI:** 10.64898/2026.09.23.753807

**Authors:** Mélanie Fourgeaud, Axel Tollance, Stéphane Koenig, Maud Frieden

## Abstract

Skeletal muscle is a dynamic tissue able to regenerate in response to injury. This regeneration relies primarily on the activation of precursor cells, known as muscle stem cells (MuSC). Upon activation, skeletal MuSC re-enter the cell cycle and proliferate as myoblasts, which subsequently either differentiate and fuse to form new fibers or return to quiescence to maintain long-term regenerative capacity. From primary human myoblasts, we generated reserve cells (RCs) – quiescent cells similar to MuSC – and investigated their activation after stimulation with a growth medium containing serum. Here, we show that Orai3 calcium channel is primarily expressed in RC, where it promotes RC activation in a calcium-independent manner. We demonstrate that Orai3 does not contribute to store-operated calcium entry (SOCE) or calcium response induced by serum stimulation in RC cells, supporting the hypothesis that Orai3 plays a unique role in RC, distinct from its calcium channel activity. Using a protein proximity assay (BioID), we identified the large scaffold protein AHNAK2 as a potential partner of Orai3. AHNAK2 is expressed at high level in myotubes, and has a similar effect on RC activation than Orai3. Both Orai3 and AHNAK2 are necessary for proper myoblast differentiation and RC activation, while acting through different signaling pathways. Specifically, we show that Orai3 directly influences the RC fate, whereas AHNAK2 facilitates RC activation via signals derived from myotubes. These findings provide new insights into the molecular mechanisms underlying the activation and quiescence of human MuSC.

## Background

Skeletal muscle function is a vital component of good health. Deterioration in this function results in mobility limitations and metabolic deficits, which harm an individual’s quality of life. Daily life use of skeletal muscle might lead to damage of various extents, which can, however, be repaired through muscle regeneration. This regeneration process is mainly driven by muscle stem cells (MuSC), which are adult stem cells actively maintained in a quiescent state [1]. Upon injury, MuSC become activated and re-enter the cell cycle [2]. The regulation of this process depends on several mechanisms, including the alteration of the MuSC microenvironment and the release of secreted factors by the various surrounding cells [3]. Activated MuSC proliferate as myoblasts, which then either differentiate and fuse to repair damaged muscle fibers, or return to quiescence to replenish the MuSC pool [4,5]. Their return to quiescence is essential for maintaining long-term repair capacity. It results from the increased activation of specific signaling pathways, such as the Notch pathway, that downregulates the myogenic factors [6,7].

In humans, the molecular mechanisms involved in the activation and self-renewal of MuSC remain poorly understood, primarily due to the challenges of monitoring these processes ex vivo. In fact, extracting MuSC from a muscle biopsy is enough to activate them and alter their gene expression [8]. Applying *in situ* fixation of mouse muscles, Machado et al. have obtained the gene expression profile of quiescent and activated murine MuSC [9]. Although this approach has been successful in describing regulators of quiescence and early activation, it is not translatable to human studies. To study human quiescent and activated MuSC, we use an in vitro model of human primary muscle cells where we recapitulate the process of regeneration. In the absence of serum, approximately 70% of myoblasts undergo differentiation and fuse to form giant polynucleated cells known as myotubes. The remaining myoblasts return to quiescence and acquire stem cell characteristics [10,11]. These stem cell-like cells, called reserve cells (RC), allow us to study both the transition from myoblasts to quiescent RC and the activation of RC following stimulation [12].

In our previous work, we used this culture model to study serum-induced RC activation. We showed that serum stimulation elicited a robust calcium (Ca^2+^) response in RC but not in myotubes. Surprisingly, we revealed that RC’s re-entry into the cell cycle was independent of the Ca^2+^ response, as preventing all Ca^2+^ fluxes did not impede RC activation. On the contrary, the migration of RC relies on Ca^2+^ signals [13]. Recently, it was reported that the Ca^2+^ channels from the Orai family could be involved in physiological processes such as proliferation, but independently of their channel function [14]. Orai channels are the main channels responsible for Ca^2+^ entry activated by store depletion, called store-operated Ca^2+^ entry (SOCE;[15]). In the present study, we investigated the putative role of Orai3 in RC activation and found that this channel is required for their activation but in a Ca^2+^-independent manner. In addition, using biotin proximity labelling for protein interaction (BioID), we uncover AHNAK2, a large scaffold protein, as a potential binding partner of Orai3. Like Orai3, AHNAK2 is implicated in RC activation, and both proteins appear necessary for proper myoblast differentiation, while acting through different signalling pathways.

## Methods

### Primary human myoblast cell culture

Human primary muscle stem cells were isolated from semitendinosus muscle samples obtained postoperatively from patients undergoing orthopedic surgery, as surgical waste. The donors had no known muscular diseases. All samples were collected anonymously after written consent and approval by the University of Geneva (protocol CCER no. PB_2016-01793 (12-259) accepted by the Swiss Regulatory Health Authorities and approved by the "Commission Cantonale d’Ethique de la Recherche" from the Geneva Cantonal Authorities, Switzerland). Muscle stem cells purification was performed as previously described [11]. The myoblasts were cultured in a growth medium (GM) until they reached confluence. Then, the GM was replaced with a differentiation medium (DM) to initiate differentiation. The composition of the GM and DM has been described previously [16] with the exception that the DM used in the present study was not supplemented with horse serum. Myoblasts were differentiated for 48h to obtain myotubes and RC. RC activation was induced by readdition of GM for 24 hours. When needed, myotubes and RC were separated by partial trypsinisation. The detached myotubes were either used to extract RNA, proteins or were discarded. The remaining RC were either subjected to a second round of trypsinization and filtered (20 µm) to obtain RNA or proteins, or left attached for activation with GM. To prepare the conditioned media used to activate RC, myoblasts transfected with siRNA were differentiated for 48h and then incubated with GM. After 24h, the culture supernatant containing the cell secretome was collected, filtered and applied to adherent RC.

### siRNA transfection

Myoblasts were transfected using Lipofectamine RNAiMAX transfection reagent (Thermo Fisher Scientific, cat. No 13778150) according to the manufacturer’s instructions using 40 pmol of siRNA. Differentiation was induced 48 hours after siRNA transfection. Two different siRNA targeting Orai3 and AHNAK2 were used to confirm the results obtained. The sequences of all siRNA are listed in Table 1.

**Table 1.**
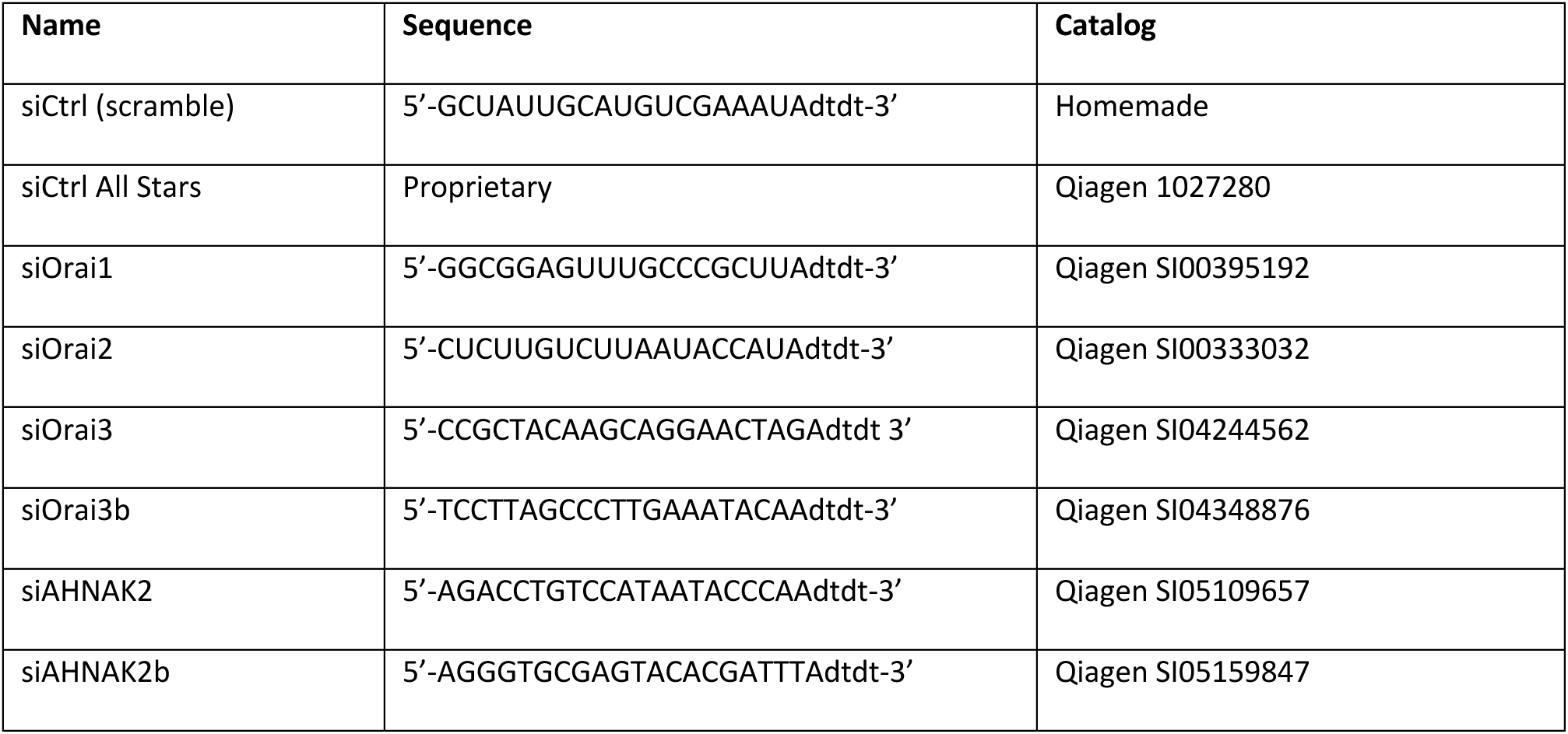
siRNA sequences.

### RNA isolation and RT-qPCR

Total RNA was extracted using TRI reagent solution (Invitrogen, cat. No AM97738) according to the manufacturer’s instructions. All following steps were performed in the iGE3 Genomics Platform of the University of Geneva. Briefly, 0.5 μg of total RNA was reverse-transcribed with the PrimeScript RT reagent kit (TAKARA, Bio Company, Japan) according to the manufacturer’s protocol. PCR was performed on an SDS 7900 HT instrument (Applied Biosystems). Raw threshold-cycle (Ct) values were obtained with SDS 2.2 (Applied Biosystems). Targeted transcript levels were normalized by the expression of two reference genes, *B2M* (β2-microglobulin) and *EEF1A1*. The primers used are listed in Table 2.

**Table 2.**
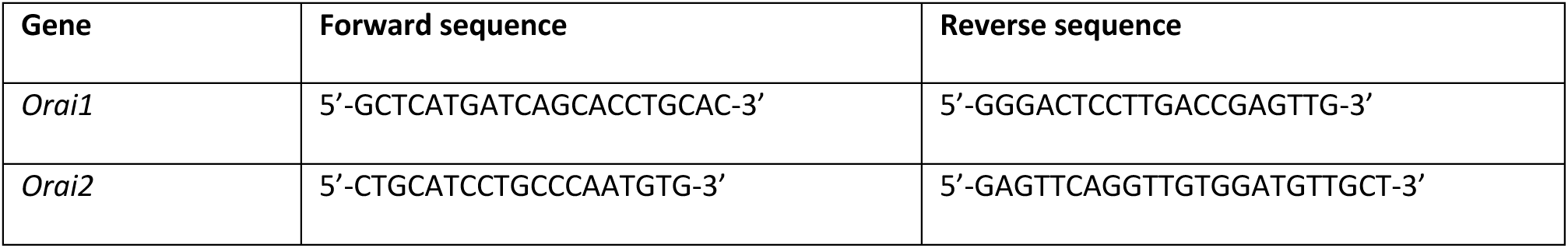
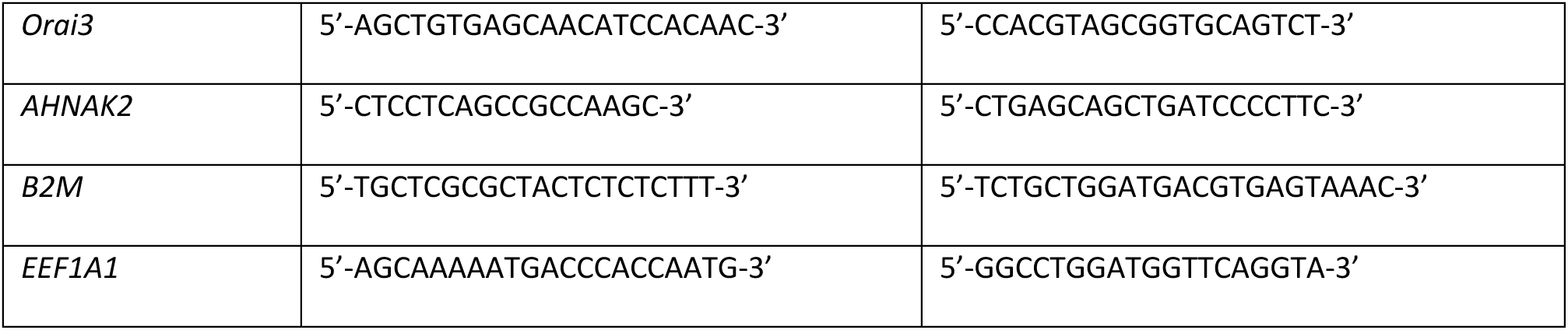
List of primers.

### Immunofluorescence

Cells were fixed in PBS with 4% paraformaldehyde for 15 minutes, followed by permeabilization and blocking in PBS containing 0.3% of Triton X-100 and 5% goat serum for 30min. Primary antibodies (Table 3) were incubated at 4°C overnight. Secondary antibodies (Table 4) were incubated for 75 minutes at room temperature. RC activation was evaluated by EdU (5-ethynyl-2’-deoxyuridine) assay (Click-iT™ Plus EdU Cell Proliferation Kit for Imaging, Alexa Fluor™ 647 dye, C10640, ThermoFisher) according to manufacturer’s instructions. Nuclei were stained using ProLong™ Glass Antifade Mountant with NucBlue™ (Hoechst 33342) (ref. P36931, Life Technologies). The acquisition of images was performed with a widefield AxioImager M2 microscope (Zeiss, Germany) through a 10X or 20X objective. The analysis of the immunofluorescence images was performed using ImageJ and Qupath softwares. To quantify cell differentiation, the threshold for MEF2C positivity was determined at 48h after differentiation using kernel density estimation in Matlab (R2024a, Bioimaging Core Facility, University of Geneva). The same threshold was applied to the time points 0h and 24h.

**Table 3.** Primary antibodies.

| Name | Catalog no | Host | Dilution |
| --- | --- | --- | --- |
| anti-α-actinin | A7811, Sigma | mouse | 1:500 |
| anti-MEF2c | 5030S, Cell Signaling | rabbit | 1:500 |

**Table 4.** Secondary antibodies.

| Name | Catalog no |
| --- | --- |
| Alexa Fluor® 488-conjugated goat anti-mouse IgG | A11029, Life Technologies |
| Alexa Fluor® 546-conjugated goat anti-rabbit IgG | A11030, Life Technologies |

### Calcium imaging

Culture of myotubes and RC after 48h of differentiation were loaded with Fura2-AM (2 µM, F1201, ThermoFisher) at RT and in the dark for 30min in a medium containing 135 mM NaCl, 5 mM KCl, 1 mM MgCl_2_, 10 mM Hepes, 2 mM CaCl_2_, 10 mM glucose, pH adjusted at 7.45 with NaOH (CA medium). Following the incubation, cells were washed and kept for 10 min in the same medium to allow the de-esterification of the dye.

Ratiometric images of Ca^2+^ signals were recorded using a Zeiss Axio Observer A1 microscope equipped with a Lambda XL illumination system (Sutter Instrument, Novato, CA, USA) and switching the excitation wavelengths between 340 nm (ET340x; Chroma) and 380 nm (ET380x; Chroma) with images taken every 2 seconds. Emission was collected through a 415 DRLP dichroic mirror and a 510WB40 emission filter (Omega Optical) by a cooled 16-bit CMOS camera (pco.Edge sCMOS, Visitron Systems, Puchheim, Germany). Image acquisition was performed with the VisiWiew software version 4.4.0.11 (Visitron Systems, Puchheim, Germany), and the analysis was done with Fiji and Matlab software.

For the fetal calf serum stimulation, cells were recorded for 2 min before stimulation and recorded for a total of 30 minutes. The stimulation was performed with CA medium + 15% Fetal calf serum. For the SOCE protocol, cells were stimulated in 250 nM free Ca^2+^ solution, containing 135 mM NaCl, 5 mM KCl, 3 mM MgCl_2_, 10 mM Hepes, 10 mM glucose, 1 mM EGTA and 0.678 mM CaCl_2_, pH 7.4 (NaOH) with 1 μM of thapsigargin for 10 min followed by 2 mM Ca^2+^ re-addition to measure SOCE. A 250 nM free Ca^2+^ was used instead of Ca^2+^-free to minimize myotube detachment. For the Mn^2+^ quench experiments, cells were excited at 360 nm (360BP10; Omega Optical), the isosbestic point of Fura-2. After 10 minutes of thapsigargin stimulation in CA medium, 500 µM Mn^2+^ was added, and the slope of the fluorescence quench was measured as an estimate of the Ca^2+^ entry.

### Biotin-Identification and protein analysis

The BioID-Orai3 fusion gene was generated by cloning the human Orai3 gene into a plasmid encoding the BioID ligase protein (Addgene #35700). Twenty-four hours after transfection, 50 µM biotin was added to the myoblast culture for an additional 24 hours. Control conditions included cultures without biotin and untransfected myoblasts treated with 50 µM biotin for 24 hours. Biotinylated proteins were immunoprecipitated using the Pierce™ MS-Compatible Magnetic IP Kit (streptavidin) (Thermo Scientific, Cat. No. 90408), following the manufacturer’s instructions.

Proteins were identified through LC-ESI-MS/MS mass spectrometry at the Proteomics Core Facility, University of Geneva. The resulting peaklist files were searched against the human Reference Proteome database (UniProt, 2018-06, containing 21044 entries) and an in-house database of common contaminants using Mascot (Matrix Science, London, UK; version 2.5.1). Trypsin was chosen as the enzyme, allowing for one missed cleavage. The Mascot results were validated with Scaffold 5.0.0 (Proteome Software).

### Statistics

Data were analyzed with GraphPad Prism and presented as mean ± error bar (SD or SEM as indicated). Statistics were performed using the test specified for each figure.

## Results

### Orai3 downregulation decreases RC activation

We first evaluated the mRNA expression levels of the three Orai proteins during the differentiation of human primary myoblasts from different donors. Proliferating myoblasts were differentiated for 48 hours to form myotubes and quiescent RC. RT-qPCR was performed on mRNA extracted from the three different cell types obtained from each population: myoblasts, myotubes, and RC. Orai1 mRNA showed the highest levels in human primary myotubes compared to myoblasts and RC, while Orai2 mRNA showed no variation among the cell types. Conversely, Orai3 expression was higher in RC than in the other muscle cell types, suggesting a possible specific role for Orai3 in RC (Figure 1A). We then assessed RC activation following Orai downregulation to determine whether these channels regulate RC’s quiescent state. Myoblasts were transfected with siRNA 48 hours before starting differentiation. After 48 hours of differentiation, RC activation was induced by GM stimulation for 24 hours, and RC cell cycle entry was tracked through EdU incorporation. The percentage of EdU-positive nuclei was measured in the RC population, identified as cells with MEF2C-negative nuclei. Neither siOrai1 nor siOrai2 significantly affected RC activation. Notably, downregulating Orai3 (siOrai3) led to a 30% reduction in EdU-positive nuclei compared to siCtrl (Figure 1B-C). qRT-PCR confirmed that the siRNAs effectively knocked down their targets without lowering other Orai isoforms (Figure S1). The use of a second siRNA targeting Orai3 (siOrai3b) further supported the role of Orai3 in RC activation (Figure 1D). Our data indicate that Orai3, which is predominantly expressed in RC, is key to activating RC.

**Figure 1.**
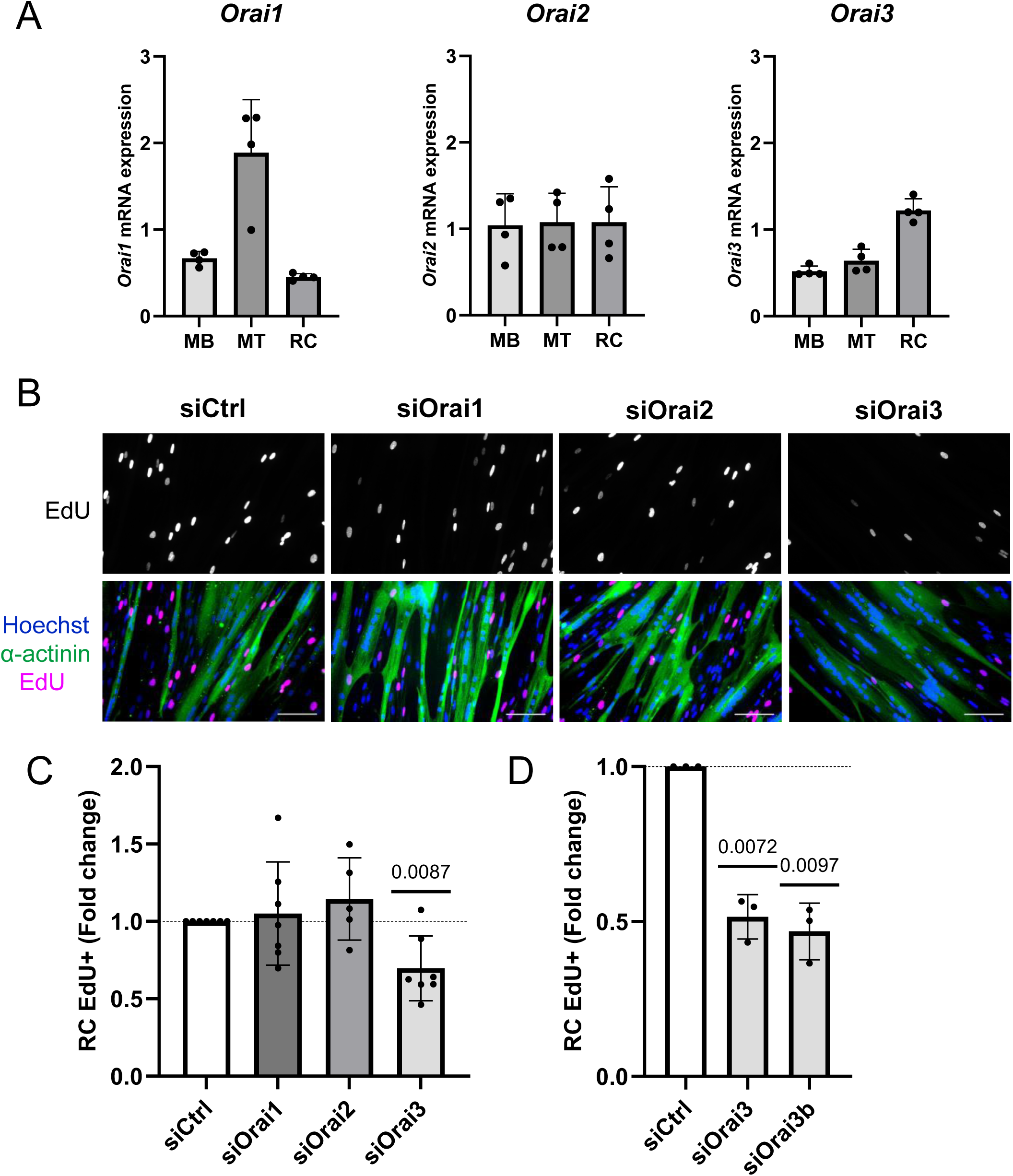
Orai3 downregulation decreases reserve cell activation. **A.** Relative quantification of Orai transcripts in human primary muscle cells. Transcript levels are normalized by the expression of reference genes (*B2M, EEF1A1*). Results are expressed as mean ± SD. N= 4 donors. **B.** Myoblasts (MB) were differentiated for 48h to obtain myotubes (MT) and reserve cells (RC). RC activation was induced by growth medium (GM) for 24h, and the re-entry into the cell cycle was assessed by EdU incorporation. Immunofluorescence images of cells transfected with siRNA after 24h of activation. Proliferative cells were stained with EdU, myotubes with α-actinin and nuclei with Hoechst. Scale bar, 100 μm. **C.** Quantification of EdU-positive RC after 24h of activation. This panel reports a total of 7 experiments representing 4 different donors. **D.** Confirmation of Orai3 effect on RC activation using a second siRNA. N=3 donors. The dotted horizontal lines represent the normalized control value. Bars show mean fold change ± SD (one sample t-test).

### Orai3 does not participate in the SOCE of RC

Since we previously demonstrated that RC activation is independent of Ca^2+^ signaling, the involvement of Orai3 in this process was unexpected. Therefore, we aimed to assess the involvement of Orai3 channels in the SOCE of RC (Figure 2A). We used Orai1 downregulation as a positive control because of its well-established role in SOCE. As expected, the amplitude and slope of SOCE significantly decreased in RC transfected with siOrai1. However, downregulating Orai3 did not cause a significant change in SOCE, neither in the amplitude nor in the slope (Figure 2B-C). We confirmed this result using the Mn^2+^ quench technique, and obtained consistent results that highlight the role of Orai1, but not Orai3, in RC’s SOCE (Figure 2D). Analysis of SOCE in myotubes surrounding the RC shows that Orai3 contributes to SOCE in these cells, while the main player remains Orai1 (Figure S2 A-B). These results demonstrate that, as expected, Orai1 is the primary isoform responsible for SOCE in RC and myotubes, whereas Orai3 only participates in SOCE in myotubes.

**Figure 2.**
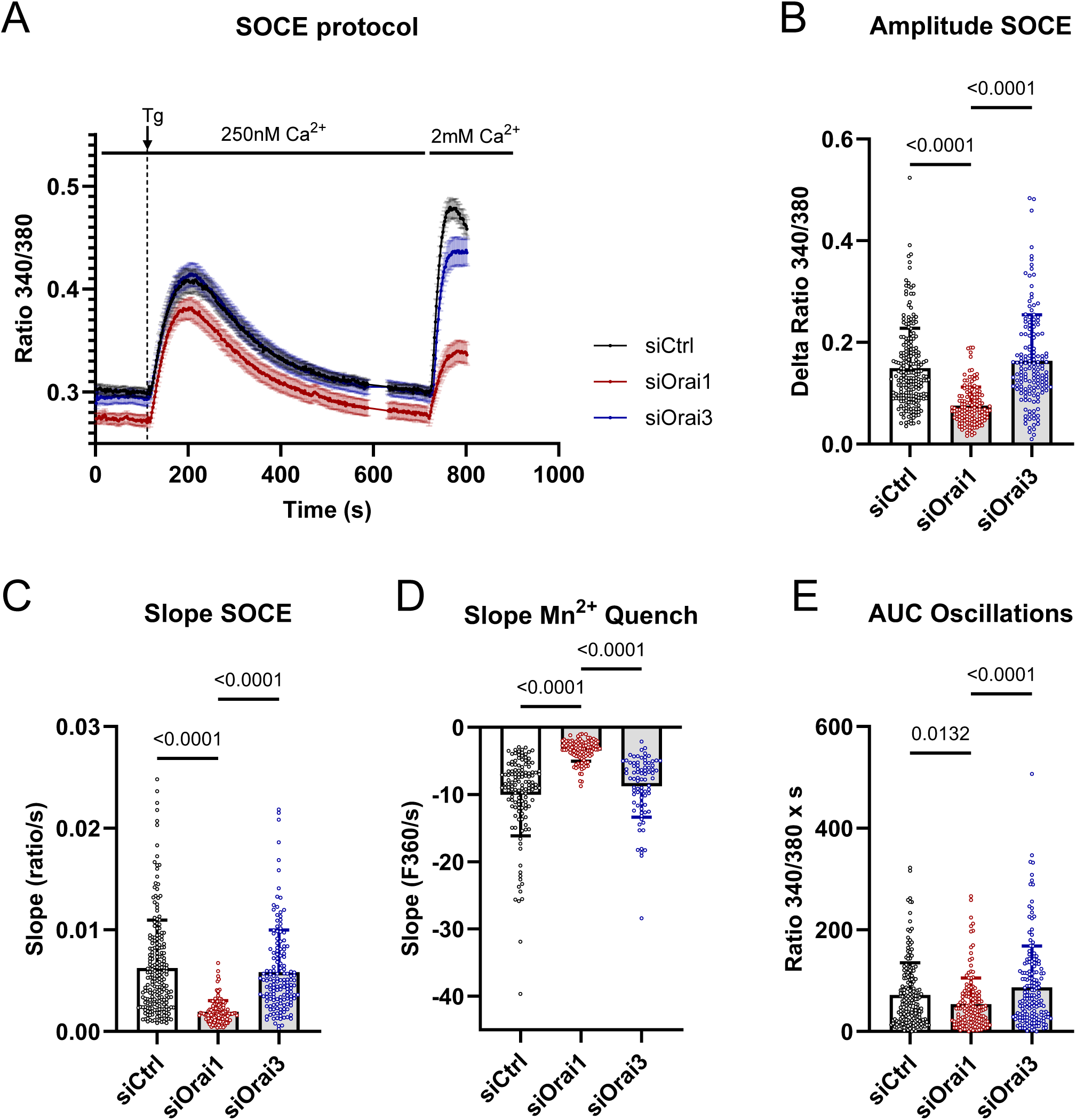
Orai3 knockdown does not impact calcium signaling in RC. Myoblasts were transfected with siCtrl, siOrai1 or siOrai3 before induction of differentiation, and the cells were loaded with Fura-2 48h after differentiation to measure cytosolic Ca^2+^ concentration (A-C, E) or the fluorescence quench (D). **A.** Store-operated Ca^2+^ entry (SOCE) protocol. The cells were maintained in a 250nM free Ca^2+^ medium and stimulated with 1μM of thapsigargin (Tg) for 10 minutes, followed by the re-addition of 2 mM Ca^2+.^ The Ca^2+^ traces show the RC mean ratios ± SEM of one representative experiment. **B-C**. Quantification of the SOCE amplitude (**B**) and slope (**C**) with siCtrl, siOrai1 and siOrai3. **D**. RC were stimulated with 1 µM Tg for 10 minutes, followed by the addition of 500 µM Mn^2+^ to assess Ca^2+^ entry. Quantification of the slope of Mn^2+^ quenching. **E**. Quantification of the area under the curve (AUC) of the Ca^2+^ response following serum stimulation. Bars show mean ± SD (Dunn’s multiple comparison test following the Kruskal-Wallis test). SOCE experiment (B-C) and AUC of calcium oscillations (E) report a total of 4 experiments representing 3 different donors (> 25 cells per condition per biological replicate). Mn^2+^ quench experiments (D) were performed on 2 donors.

Our investigation then focused on the role of Orai3 in physiological Ca^2+^ signaling, such as the oscillatory response triggered by serum stimulation [13]. To determine Orai’s impact, we measured the area under the curve (AUC) of the Ca^2+^ oscillations during 25 minutes after downregulating Orai1 or Orai3. Results showed that downregulating Orai1 reduces Ca^2+^ signaling induced by serum, whereas downregulating Orai3 has no effect (Figure 2E). These findings confirm that, although Orai3 expression is higher in RC than in myotubes, it does not function as a Ca^2+^ channel in RC. Instead, it plays a different role, particularly in activating the RC, which is likely independent of its Ca^2+^ channel activity.

### AHNAK2 is a new potential partner of Orai3

To understand the role of Orai3 in RC activation, we used the BioID technique to identify potential Orai3 protein partners in human primary myoblasts. We created a plasmid encoding a fusion protein of Orai3 and the BirA enzyme at the N-terminus. The BirA enzyme covalently attaches biotin to lysine residues of nearby target proteins within approximately 10 nm [17]. After purification with streptavidin beads, we identified the biotinylated proteins via mass spectrometry. Among the identified proteins, we kept only those that did not appear in control conditions (BioID-Orai3 without biotin or biotin alone). STIM1 and STIM2 were part of the Orai3 protein complex, while Orai1 and Orai2 were not detected (Table S1). Orai3 proteins were found among the labelled proteins, indicating that Orai3 forms homomultimers. Among the other proteins identified, we selected AHNAK2 for further study because it is a scaffold protein found in skeletal muscles and was reported to interact with ion channels in cardiomyocytes [18]. However, nothing is known regarding its role in MuSC.

### AHNAK2 exhibits similar effects on RC activation as Orai3

Western Blot analysis revealed that AHNAK2 is expressed in the RC and is strongly present in the myotubes (Figure S3). To investigate the function of AHNAK2 in regulating RC fate, we silenced its expression with siRNA, similar to our approach for Orai3. The silencing of AHNAK2 did not influence Orai3 mRNA levels, and the reverse was also true (Figure S4A). Following GM stimulation, knocking down AHNAK2 reduced the percentage of EdU-positive RC by about 60%, indicating decreased RC activation. This result was confirmed with a second siRNA targeting AHNAK2 (Figure 3A-B). Additional experiments assessed the effects of silencing both AHNAK2 and Orai3 together or separately on the same cell batches (Figure 3C). The data showed that both siOrai3 and siAHNAK2 similarly affect RC activation, with no additive effect when combined. RT-qPCR verified the effective reduction of AHNAK2 and Orai3 expression by siRNAs, whether used alone or together (Figure S4B). Overall, these results show that AHNAK2, like Orai3, is involved in RC activation.

**Figure 3.**
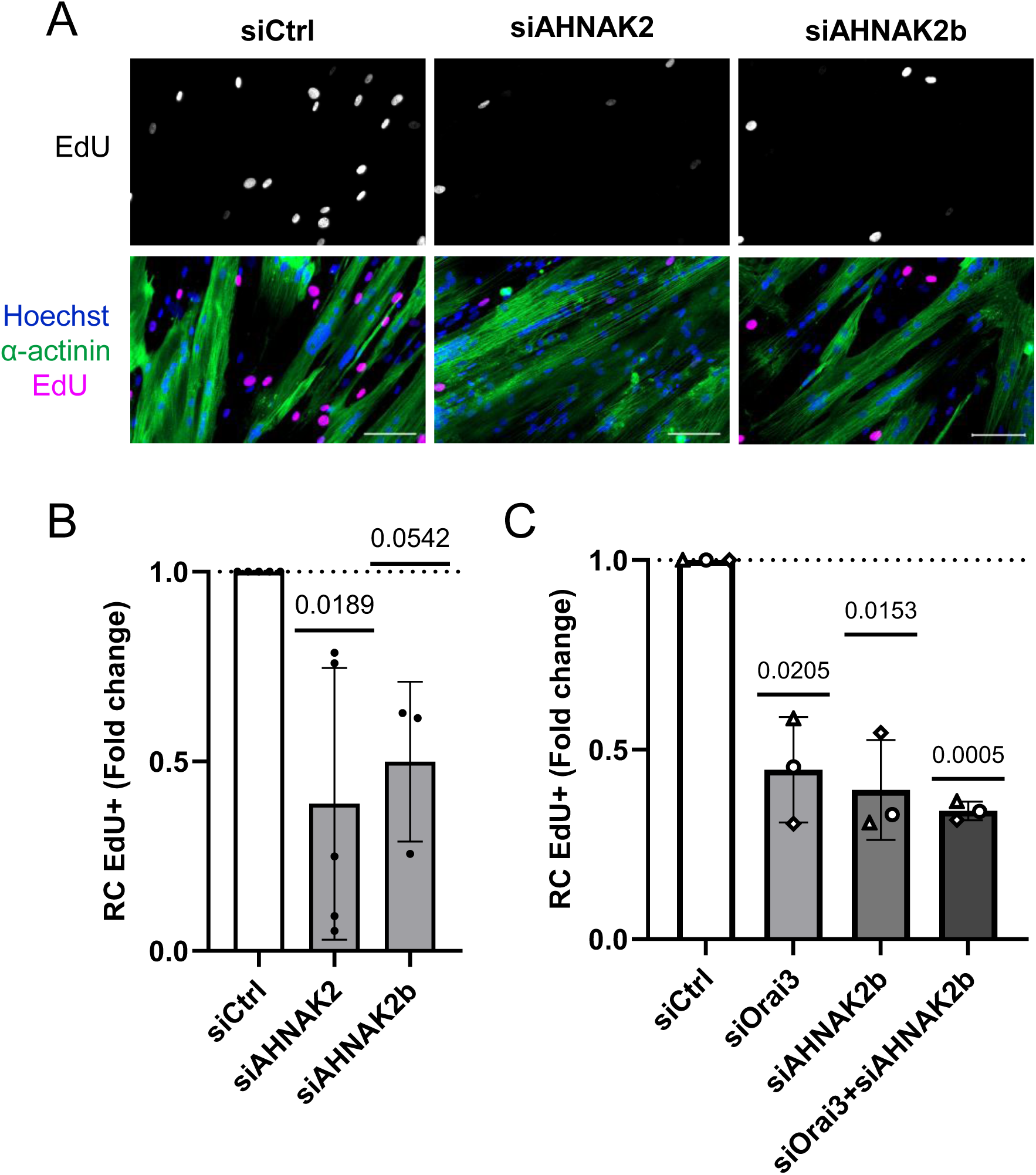
AHNAK2 downregulation decreases reserve cell activation. Myoblasts (MB) were differentiated for 48h to obtain myotubes (MT) and reserve cells (RC). RC activation was induced by growth medium (GM) for 24h, and the re-entry into the cell cycle was assessed by EdU incorporation. **A.** Immunofluorescence images of cells transfected with siRNA after 24h of activation. Proliferative cells were stained with EdU, myotubes with α-actinin and nuclei with Hoechst. Scale bar, 100 μm. **B-C.** Quantification of EdU-positive RC after 24h of activation. The dotted horizontal lines represent the normalized control value. Bars show mean fold change ± SD. N=5 (B) and 3 (C) independent donors (one sample t-test).

### Orai3 or AHNAK2 depletion affects commitment toward myotubes

The decrease of RC activation in conditions of siOrai3 or siAHNAK2 might be the consequence of an alteration in the RC quiescent state. In line, a change in quiescence could be linked to a shift in myoblasts’ commitment to differentiation. We thus examined whether depleting Orai3 and AHNAK2 before differentiation would affect the differentiation process. To study this, we evaluated the differentiation rate after one and two days of differentiation following the downregulation of Orai3 or AHNAK2. Alpha-actinin and MEF2C expression was assessed by immunofluorescence to follow cell differentiation (Figure 4A). The alpha-actinin staining showed that the total myotube surface enlarged after AHNAK2 knockdown (94’273 ± 37’123 μm^2^ in siCtrl vs 140’298 ± 17’057 μm^2^ in siAHNAK2), whereas transfection with siOrai3 did not impact myotube size after 48 hours of differentiation (Figure 4B). Reducing Orai3 levels resulted in more MEF2C-positive nuclei after 48 hours in differentiation medium (67.0 ± 3.1 % in siCtrl vs 76.7 ± 2.5% in siOrai3). A trend towards more MEF2C-positive nuclei can already be observed in myoblasts 48 hours post-siOrai3 transfection (T0). In contrast, downregulating AHNAK2 delayed differentiation, with a 18% drop in MEF2C at 24 hours, but no difference at 48 hours was observed (Figure 4C). This indicates that the increase in the surface area of myotubes downregulating AHNAK2 is not linked to an increase in the fusion index, which is estimated by calculating the percentage of MEF2C-positive nuclei. Our results indicate that downregulating AHNAK2 affects myotubes, evidenced by a slowdown in the differentiation process and resulting in larger myotubes. These findings indicate that while both Orai3 and AHNAK2 are involved in RC activation, they have distinct roles during the differentiation process.

**Figure 4.**
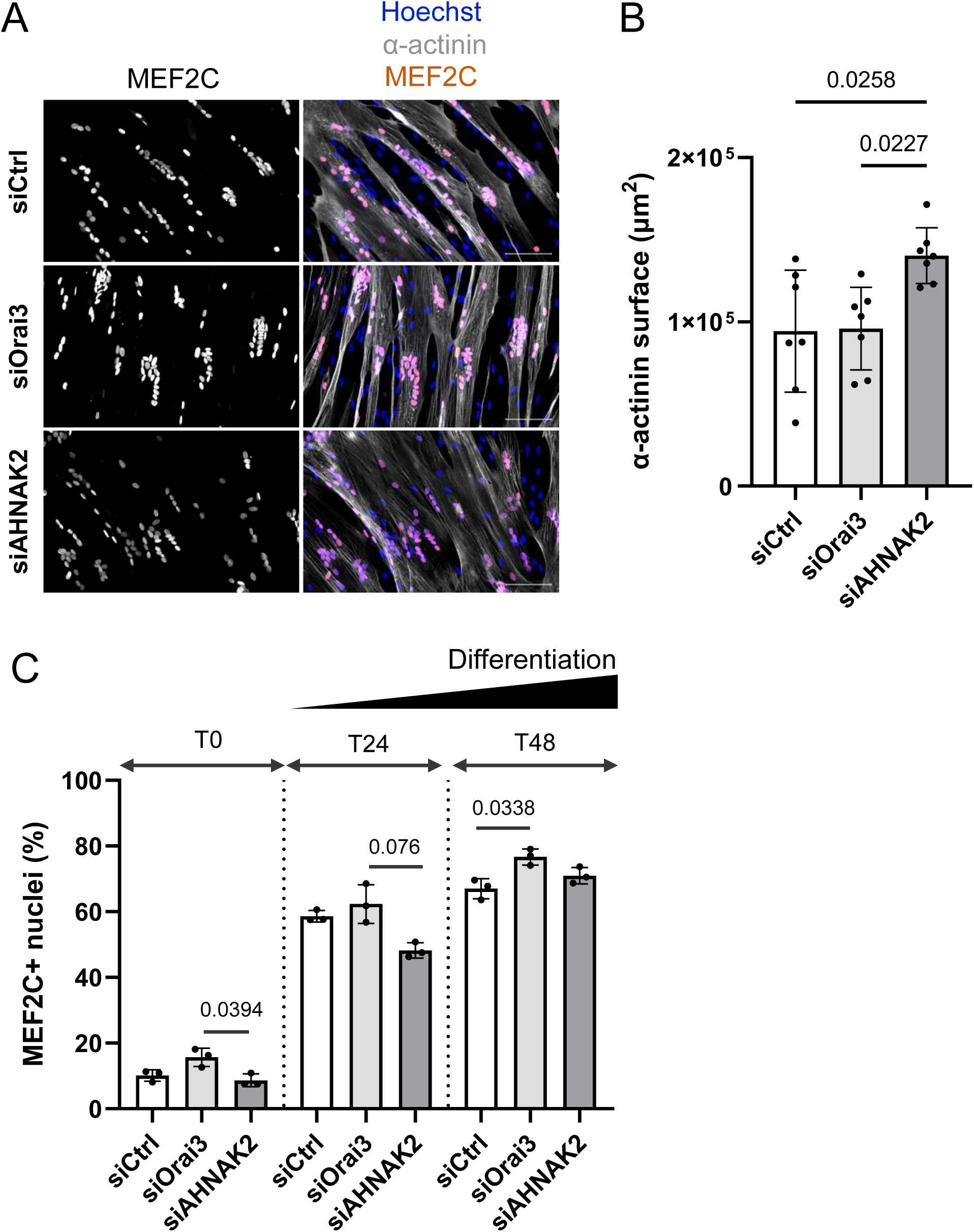
Depletion of Orai3 or AHNAK2 affects myotube differentiation. **A.** Immunofluorescence images of myoblasts transfected with siCtrl, siOrai3 or siAHNAK2 after 48 hours of differentiation. Differentiated cells were stained with MEF2C, myotubes with α-actinin and nuclei with Hoechst. Scale bar = 100 μm. **B.** Quantification of α-actinin surface after 48h of differentiation. N=7 donors. **C.** Quantification of MEF2C positive nuclei (MEF2C+) performed at 0, 24h, or 48h after induction of differentiation. N=3 donors. Bars show mean ± SD (Dunn’s multiple comparison test following the Kruskal-Wallis test).

### AHNAK2 affects RC activation only in the presence of myotubes

Reducing Orai3 levels leads to more nuclei positive for MEF2C, suggesting a decline in commitment to the RC pathway, where Orai3 is more expressed. Conversely, the downregulation of AHNAK2, which is more expressed in myotubes, appears to perturb myotube formation. Given the close connection between myotubes and RC pools, we tested whether the decrease in RC activation observed during the downregulation of Orai3 or AHNAK2 resulted from a change in the RC’s ability to activate or whether the changes could be attributed to a modification of myotube properties. To evaluate the potential role of myotubes in reducing RC activation after depletion of Orai3 or AHNAK2, we compared the rate of GM-induced RC activation with and without myotubes, which were removed just before GM addition (Figure 5A-C). In the presence of myotubes, as previously shown, downregulation of Orai3 or AHNAK2 leads to decreased RC activation (39 and 75 % decrease, respectively). However, when RC activation occurs immediately after removing the myotubes, the outcome diverged between siOrai3 and siAHNAK2. The reduction in RC activation caused by Orai3 downregulation persists in the absence of myotubes, indicating a failure of intrinsic capacity of RC to activate. In contrast, for AHNAK2 downregulation, removing the myotubes before activation fully restores the RC’s ability to activate. Therefore, in the case of AHNAK2 downregulation, the RC activation defect is likely caused by a modification induced by the myotubes, while the RC keep their ability to activate. RC activation by myotubes can occur via juxtacrine or paracrine mechanisms. To determine whether reduced AHNAK2 levels induce secretion of a soluble factor that inhibits RC activation, we treated RC with conditioned media from cells transfected with either control siRNA or siRNA targeting AHNAK2 (Figure 5D). As shown in Figure 5E, none of the conditioned GM reduced RC activation. These results suggest that the inhibitory effect of AHNAK2 downregulation in myotubes is not mediated by secreted factors, but rather occurs through a juxtacrine mechanism that remains to be identified. Hence, our results indicate that Orai3 and AHNAK2 are involved in RC activation through different mechanisms.

**Figure 5.**
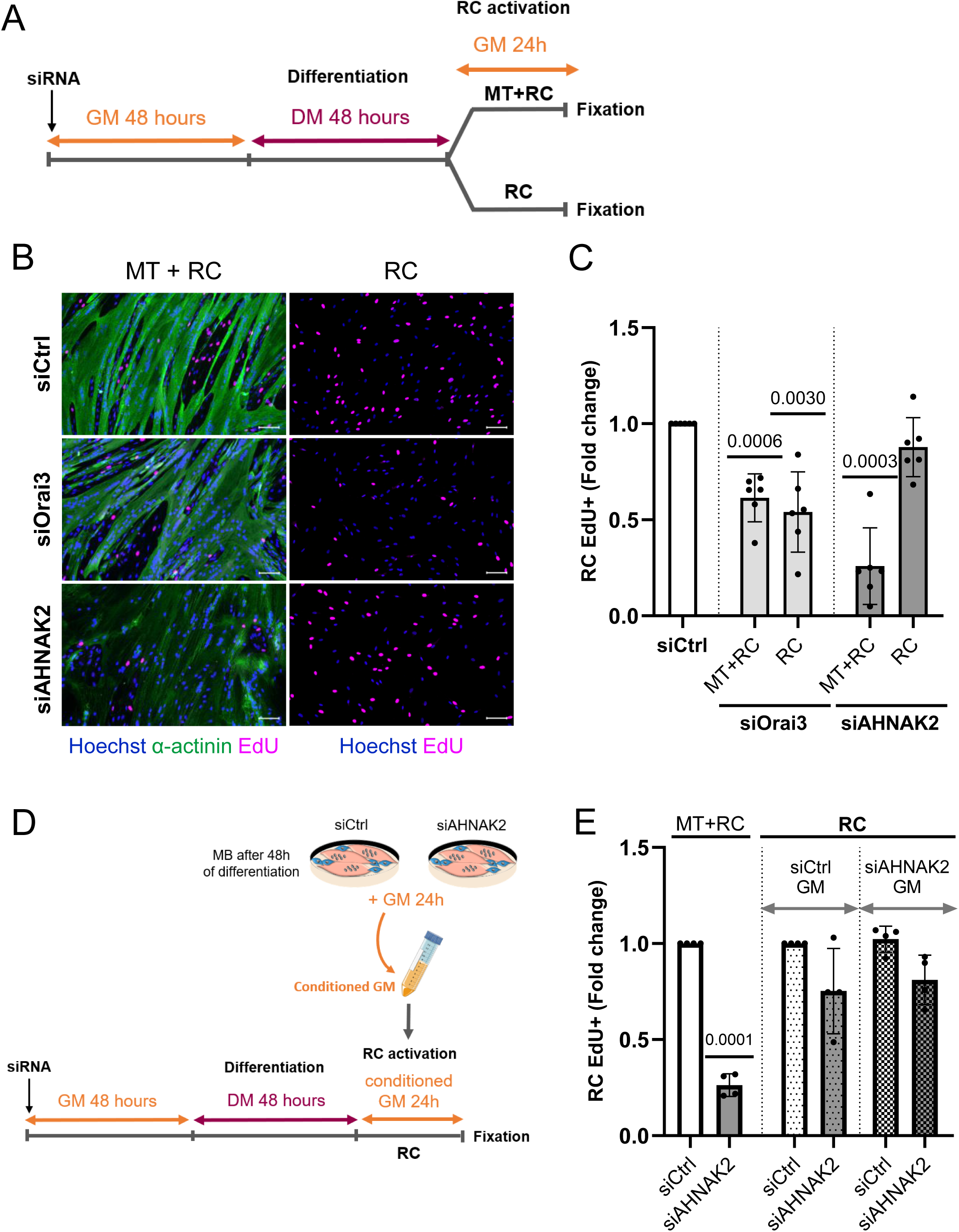
AHNAK2 decreases reserve cell activation only in the presence of myotubes. Myoblasts were differentiated for 48h to obtain myotubes (MT) and reserve cells (RC). RC activation was induced by growth medium (GM) for 24h, and the re-entry into the cell cycle was assessed by EdU incorporation, in the presence or absence of myotubes (removed by partial trypsinization). **A.** Experimental protocol is depicted. **B.** Immunofluorescence images of cells transfected with siRNA after 24h of activation in the presence (left panel) or absence of MT (right panel). Proliferative cells were stained with EdU, myotubes with α-actinin and nuclei with Hoechst. Scale bar = 100 μm. **C.** Quantification of EdU-positive RC after 24h of activation in the presence or absence of myotubes. Bars show mean fold change ± SD, N=6 donors (one sample t-test). **D.** Experimental protocol using conditioned GM containing the secretome of control myotubes or AHNAK2-depleted. **E.** Quantification of EdU-positive RC after 24h of activation with conditioned GM. Bars show mean fold change ± SD, N=4 donors (one sample t-test). GM, Growth medium; DM, Differentiation medium.

## Discussion

Identifying the mechanisms behind quiescence and activation of MuSC remains a major challenge in regenerative medicine. Although notable progress has been achieved, most studies have been performed on animal models, and their results are not always applicable to human physiology [19,20].

In this study we investigated the activation of RC using an *in vitro* model of primary human muscle cells derived from MuSC. We identified two proteins, the Ca^2+^ channel Orai3 and the scaffold protein AHNAK2, as facilitators of RC activation. Orai3 has not yet been linked to stem cell activation, and most of our knowledge on Orai3 comes from cancer research. Indeed, in many cancer cells Orai3 is overexpressed and influences cell cycle progression through multiple molecular pathways that vary with cellular context. For instance, in breast [21], lung [22], pancreatic [23], and renal cancers [24], Orai3 primarily regulates the G1/S checkpoint by modulating cyclin D1, cyclin E, CDK2, and CDK4 levels [21]. In oral/oropharyngeal squamous cell carcinoma (OSCC), Orai3 levels are higher in the cancer stem-like cell (CSC) population. Exogenous Orai3 overexpression in immortalized non-tumor oral epithelial cells enhanced SOCE and promoted malignant growth and CSC traits. Conversely, reducing Orai3 expression in OSCC cells diminished the CSC phenotype, highlighting Orai3’s key role in regulating stemness, at least in some cancer types [25]. Most of the effects linked to Orai3 overexpression in cancer cells have been proposed to occur through an enhanced SOCE that triggers downstream signalling pathways such as Akt [22] and MAP kinase/ERK1/2 [26], and/or activates transcription factors, including c-myc [26], NFATc1 [25], and HIF1α [27]. In basal breast cancer cell lines, Orai3 is involved in basal Ca^2+^ entry but not in SOCE and acts on cell migration by both Ca^2+^-dependent (through calpain) and Ca^2+^-independent (interaction with focal adhesion kinase) pathways [28]. In primary cultures of patients-derived colorectal cancer (CRC) cells, Orai3 is part of SOCE but does not contribute to cell proliferation/migration, whereas in a CRC cell line, SOCE contributes to proliferation/migration [29]. Additionally, in prostate cancer, Orai3 forms heteromers with Orai1 to create channels that promote proliferation in a store-independent manner [30]. In 2014, Borowiec et al., reported that in three different transformed cell lines, Orai1 and Orai3 are required for cell proliferation but in a Ca^2+^-independent manner, suggesting non-canonical signaling mechanisms of the channels [14]. Finally, in contexts other than cancer, Orai3 was frequently reported to exert a negative impact on SOCE [31,32]. Recently, it was also demonstrated using an Orai3^-/-^ mouse model that this channel is not implicated in T and B cells’ SOCE, even if well expressed [33]. Hence, contrary to Orai1, which is the prototypical SOCE channel, the involvement of Orai3 in SOCE depends on the cell type and the pathophysiological conditions.

Previously, we reported that RC activation happens independently of Ca^2+^ signals [13]. Notably, we showed that downregulating Orai3 levels does not impact neither SOCE in RC nor the serum-induced Ca^2+^ oscillations during RC activation. Nonetheless, we propose that Orai3 facilitates RC activation through mechanisms distinct from its Ca^2+^ channel activity, as observed in other models (see above). Although Orai3 does not participate in SOCE in RC, it contributes to SOCE in myotubes. Currently, the mechanisms underlying Orai3’s role in SOCE across different cellular environments are poorly understood, but its specific function in RC might involve unique post-translational modifications or cellular localizations beyond the plasma membrane.

Because Orai3 plays a role in RC activation independently of its Ca^2+^ channel function, one of our hypotheses was that Orai3 is part of a protein complex that enables activation of signaling pathways involved in RC activation. To identify Orai3 partners, we used biotin-proximity-dependent labeling (BioID) in myoblasts. Using this method, we identified several proteins, and we focused on AHNAK2 as a potential Orai3 binding partner for two main reasons: AHNAK2 is a giant structural protein that acts as a scaffold, and it was reported to interact with plasma membrane ion channels [18]. In addition, its homolog, AHNAK, is involved in various biological processes, including the regulation of voltage-gated Ca^2+^ channels in cardiomyocytes [18,34] and neurons [35]. Our findings indicate that AHNAK2 is highly expressed in human myotubes, and to a lesser extent in myoblasts and RC. AHNAK2 is part of the costameric network in mouse skeletal muscle, where it colocalizes with vinculin and α-actinin [36], and its C-terminal domain interacts with dysferlin and myoferlin [37]. Currently, there is no data on AHNAK2’ s role in MuSC. However, like Orai3, AHNAK2 is involved in cell proliferation and has been identified as an oncogenic factor in various cancers, consistently promoting tumor growth in pancreatic [38], lung [39] and thyroid [40], among others. In many of these models, reducing AHNAK2 levels inhibited cancer cell proliferation. Notably, AHNAK2 exhibits remarkable interaction and signaling profiles, as evidenced by cancer research. Indeed, studies of cancer cells reported that AHHAK2 interacts with proteins belonging to signaling pathways involved in the exit from quiescence of muscle stem cells. Indeed, AHNAK2 stabilizes c-MET to activate HGF/c-MET signaling [38], stimulates the MAPK pathway through effects on p-MEK, p-ERK, and p-P90RSK [39], boosts PI3K/AKT signaling [41,42], and impacts NF-κB/MMP-9 [43] and Wnt/β-catenin pathways [40].

Although no function of AHNAK2 has been described in MuSC or as a partner of Orai3 channels, either protein downregulation diminished RC activation. Given the proximity of approximately 10 nm between Orai3 and AHNAK2, based on BioID data [17,44], and their common role in RC activation, we initially hypothesized that AHNAK2 functions as a connecting factor between Orai3 and the pathways involved in RC quiescence or activation. To test this hypothesis, we downregulated both proteins simultaneously but found no additive effects on RC activation. In line, we were unable to co-immunoprecipitate endogenous AHNAK2 with Orai3-FLAG (data not shown), thereby failing to confirm a direct interaction. Furthermore, RC activation experiments with or without myotubes led to divergent outputs: RC activation was similarly impaired after Orai3 knockdown, in the presence or absence of myotubes. In siAHNAK2 condition, however, no more defects were noticed in the absence of myotubes. This demonstrated that Orai3 directly affects RC fate by acting on RC themselves, whereas AHNAK2 likely influences myotubes, which in turn affects RC activation. Factors secreted by myotubes that exert paracrine effects on RC may thus be involved. For instance, a recent study found that changes in the myofiber secretome after denervation impact MuSC activation [45]. Moreover, exosome-like vesicles released by myotubes may interact with RC, potentially altering their activation status [46]. Our results using conditioned medium are less in favor of secreted factors from myotubes. More likely, a direct interaction between myotubes and RC would explain how AHNAK2 influences RC activation. Notch signaling is a key regulator of RC quiescence, and Notch receptors and ligands are found on cell surfaces [47,48], which makes this pathway a promising candidate that remains to be tested.

We also examined how reducing Orai3 and AHNAK2 levels affects differentiation, as this could alter the state in which RC are found after differentiation and thus their activation capacity. Lowering Orai3 in proliferating myoblasts before inducing differentiation results in a higher percentage of cells expressing MEF2C during proliferation. This trend continues after 48 hours of differentiation, with more MEF2C-positive nuclei, suggesting a reduction in the RC population. The ability of activated MuSC to either self-renew or form myotubes during muscle regeneration may rely on the heterogeneity within the MuSC population, especially regarding Notch receptor and ligand expression [47–50]. Therefore, changes in the expression of Notch receptors, particularly Notch2 or Notch3, and ligands such as Dll1 or Dll4 could explain the commitment to differentiating cells at the expense of self-renewal observed when Orai3 is reduced. Conversely, downregulating AHNAK2 initially appears to slow myoblast commitment to differentiation. More significantly, it causes myotubes to enlarge, indicating a role for AHNAK2 in preserving cell structure, possibly via interactions with cytoskeletal proteins. Taken together, these data also confirm that during myoblast differentiation into myotubes and RC, Orai3 and AHNAK2 act through distinct pathways. Our findings demonstrate that Orai3 and AHNAK2 are part of the machinery involved in RC activation. We initially believed that Orai3 and AHNAK2 influenced RC activation via a common pathway. However, our data clearly indicate that Orai3 directly affects RC activation, whereas AHNAK2 promotes it via signals arising from the myotubes. Recent studies highlight the involvement of Orai3 and AHNAK2 in bestowing stem cell-like properties to cancer cells [25,51]. Notably, our results suggest that decreasing these proteins could disrupt the quiescent state and hinder re-entry into the cell cycle. These findings reveal new pathways governing cellular quiescence and may extend beyond muscle regeneration, potentially offering new targets for cancer stem cells implicated in metastasis and relapse [52].

## List of abbreviations

AUC: Area Under the Curve
CRC: Colorectal Cancer
CSC: Cancer Stem-like Cell
DM: Differentiation Medium
GM: Growth Medium
MEF2C: Myocyte Enhancer Factor 2 family
MuSC: Muscle Stem Cell
OSCC: Oral/oropharyngeal Squamous Cell Carcinoma
RC: Reserve Cell
RT-qPCR: Reverse Transcription quantitative Polymerase Chain Reaction
SOCE: Store-Operated Calcium Entry
STIM: Stromal Interaction Molecule

## Declarations

### Ethics approval and consent to participate

The University of Geneva approved this study under the protocol CCER No PB_2016-01793 (12-259).

### Consent for publication

Not applicable.

### Availability of data and materials

The datasets used during the current study are available in Yareta repository.

### Authors’ contributions

M.Fo. and A.T. performed the experiments, analyzed the data and were major contributors in writing the manuscript. S.K. and M.Fr. designed and coordinated the study and wrote the manuscript.

All authors read and approved the final manuscript.

## Acknowledgements

We thank Olivier Dupont for the excellent technical assistance, and Nicolas Liaudet for support with image analysis.

## Supplementary Figures

**Figure S1.**
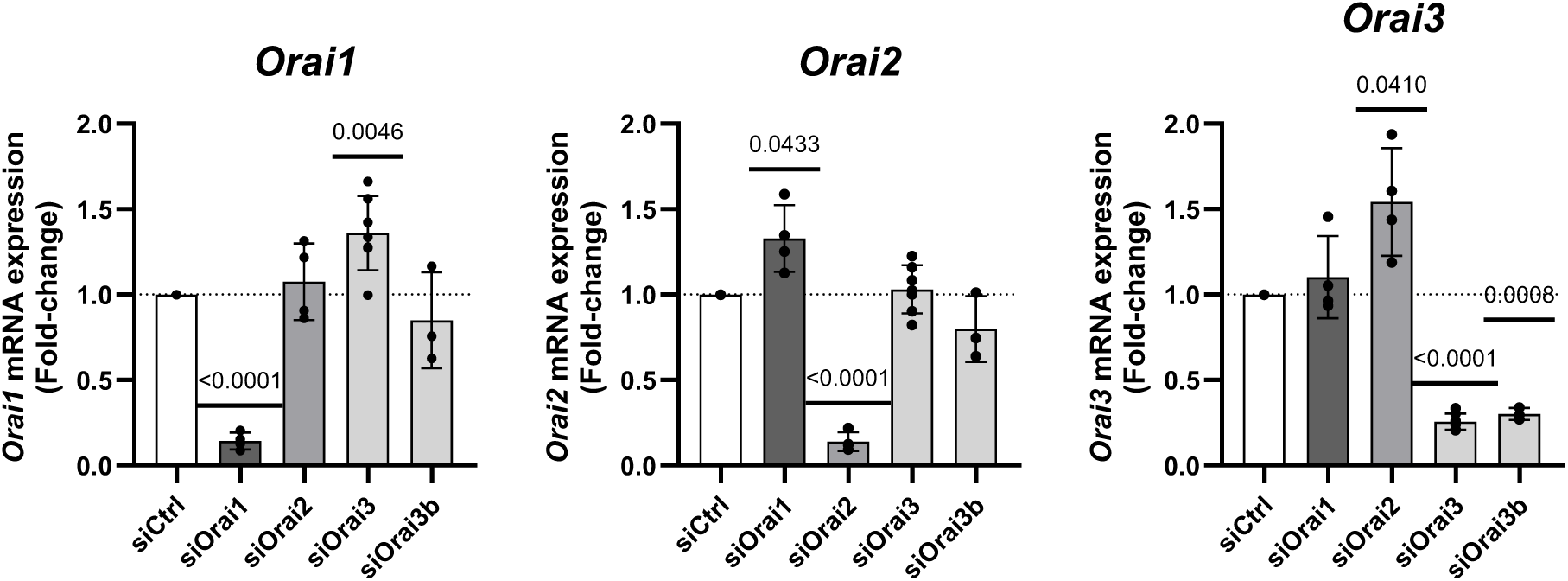
Efficiency of siRNA-mediated knockdown of *Orai* genes. *Orai* gene expression was quantified after siRNA transfections. Transcript levels are normalized by the expression of reference genes (*B2M*, *EEF1A1*). The dotted horizontal lines represent the normalized control value. Results are expressed as mean fold change ± SD. N=6 donors (one sample t-test).

**Figure S2.**
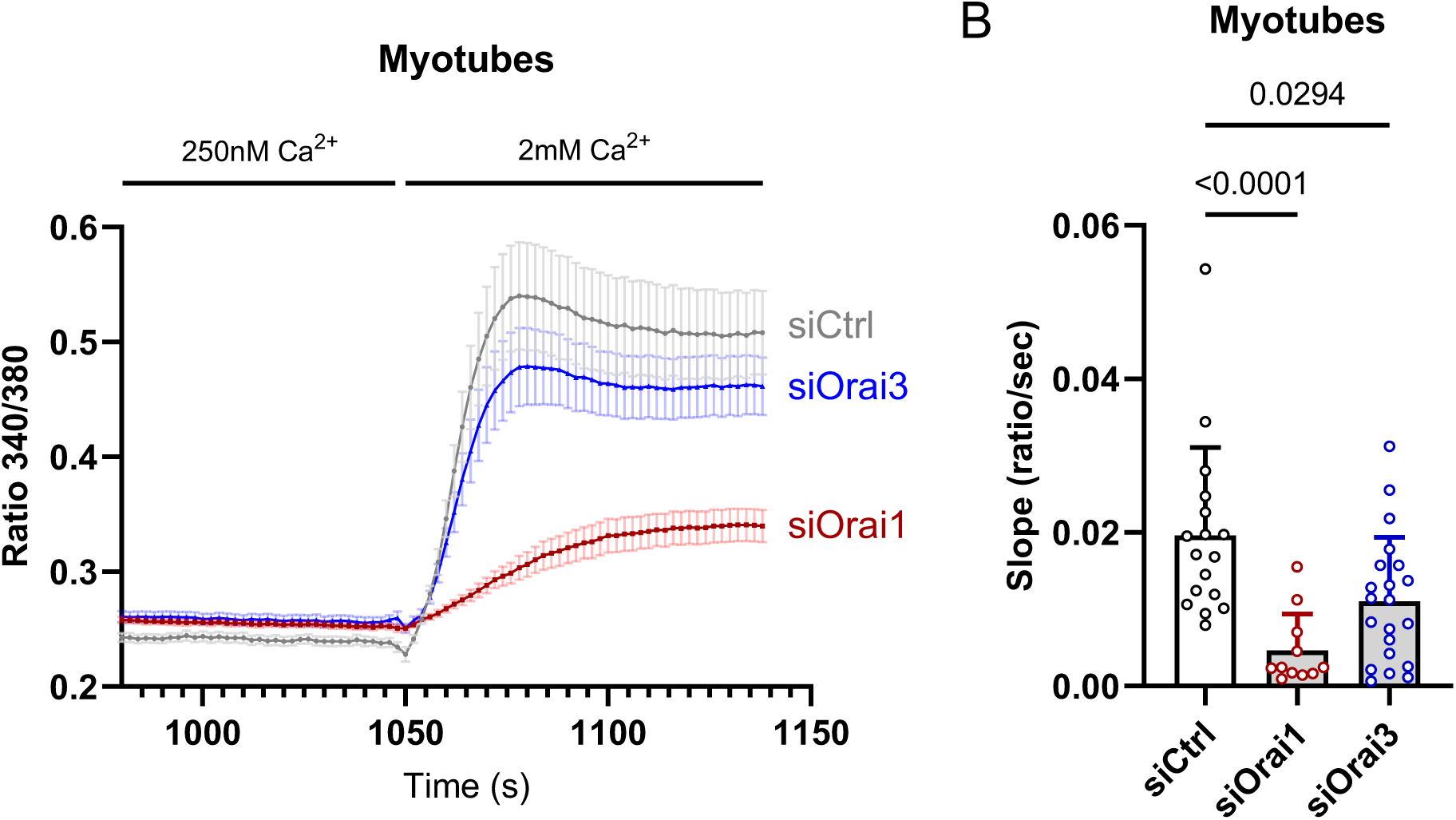
Orai3 mediates SOCE in myotubes. **A.** Store-operated Ca^2+^ entry (SOCE) traces in myotubes with siCtrl, siOrai1 or siOrai3. The cells loaded with Fura-2 were maintained in a 250nM free Ca^2+^ medium and stimulated with 1μM of thapsigargin for 10 minutes, followed by the re-addition of 2 mM Ca^2+^. Only the Ca^2+^ re-addition phase is presented, showing the mean ratio ± SEM. **B.** Quantification of the SOCE slope. Results are expressed as mean ± SD (Dunn’s multiple comparison test following the Kruskal-Wallis test). This figure reports a total of 4 experiments representing 3 different donors (n ≥ 9 myotubes per condition).

**Table S1.** List of proteins identified by BioID. Only proteins that did not have hits under control conditions (BioID-Orai3 without biotin or biotin without BioID-Orai3) are shown. Proteins are listed in alphabetical order.

| Accession code | Alternate ID | Protein name | Size |
| --- | --- | --- | --- |
| Q8IVF2 | AHNAK2 | AHNAK2 | 617 kDa |
| P06709 | birA | Bifunctional ligase/repressor BirA | 35 kDa |
| O43491 | EPB41L2 | Band 4.1-like protein 2 | 113 kDa |
| P19823 | ITIH2 | Inter-alpha-trypsin inhibitor heavy chain H2 | 106 kDa |
| P02788 | LTF | Lactotransferrin | 78 kDa |
| P43121 | MCAM | Melanoma cell adhesion molecule | 72 kDa |
| Q9BRQ5 | ORAI3 | Orai-3 | 31 kDa |
| Q9UHD8 | SEPTIN9 | Septin-9 | 65 kDa |
| O00161 | SNAP23 | Synaptosomal-associated protein 23 | 23 kDa |
| Q13586 | STIM1 | Stromal interaction molecule 1 | 77 kDa |
| Q9P246 | STIM2 | Stromal interaction molecule 2 | 84 kDa |
| O43396 | TXNL1 | Thioredoxin-like protein 1 | 32 kDa |
| O15498 | YKT6 | Synaptobrevin homolog YKT6 | 22 kDa |
| Q7Z2W4 | ZC3HAV1 | Zinc finger CCCH-type antiviral protein 1 | 101 kDa |

**Figure S3.**
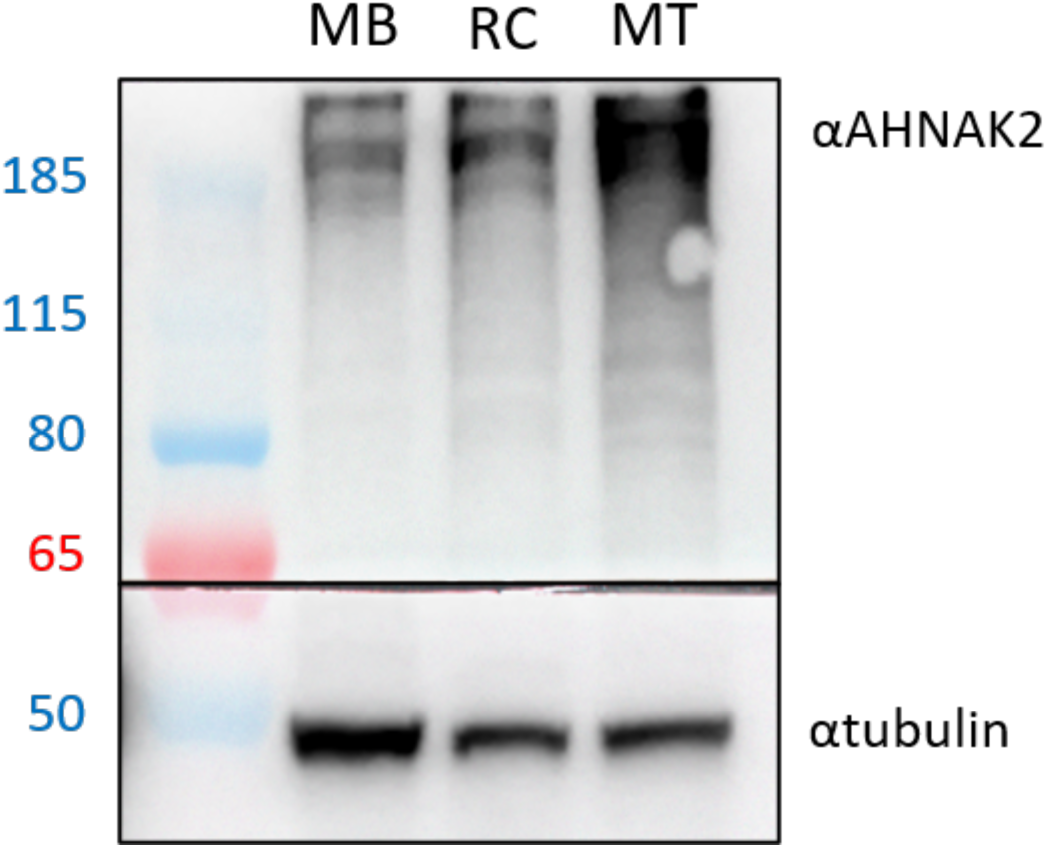
AHNAK2 protein expression in primary myoblasts (MB), reserve cells (RC) and myotubes (MT) Representative Western blot showing the protein expression of AHNAK2 in the different cell types. MB were differentiated for 48h to obtain MT and RC. The cells were lysed in a 1% CHAPS solution. Proteins were separated on 4-20% polyacrylamide gel and transferred on PVDF membranes. Membranes were blocked with a 1X PVA in TTBS for 5 min and incubated with AHNAK2 primary antibody (1/600, PA5-51781, Invitrogen, ThermoFisher Scientific) in a 5% bovine serum albumin (BSA) – TTBS solution at 4°C overnight. Blots were then incubated for 1 hour with HRP-conjugated secondary antibodies and revealed with SuperSignal West Pico PLUS Chemiluminescent Substrate (ThermoFisher Scientific) on a Fusion FX Imaging System (Vilber). Alpha-tubulin (1/10000, T9026, Sigma Aldrich) was used as a housekeeping protein.

**Figure S4.**
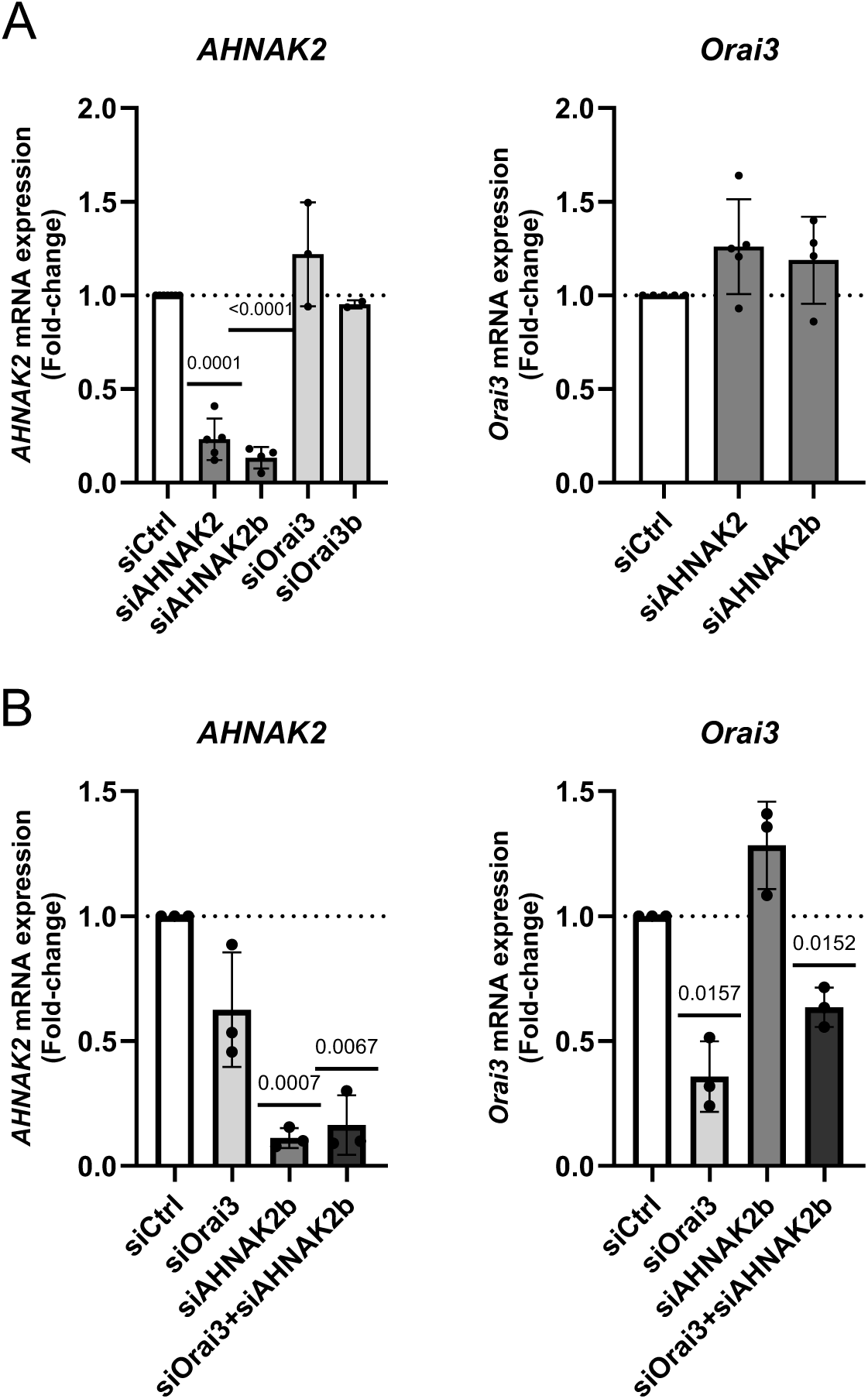
Efficiency and specificity of siRNA-mediated knockdown of *Orai3* and *AHNAK2* genes. **A-B**. Transcript levels of genes encoding for AHNAK2 and Orai3 proteins were quantified after siRNA transfection. Transcript levels are normalized by the expression of reference genes (*B2M*, *EEF1A1*). Results are expressed as mean fold change ± SD (one sample t test). The dotted horizontal lines represent the normalized control value. Panel **B** is associated with Figure 3**, panel C**.

